# The effect of E-liquid exposure on *Caenorhabditis elegans*

**DOI:** 10.1101/2020.09.14.295790

**Authors:** Ying Wang, Thomas L Ingram, Sophie Marshall, Freya Shephard, Lisa Chakrabarti

## Abstract

E-cigarettes are being promoted as a less harmful alternative to smoking tobacco. However, vaping is a new phenomenon and safety profiles have not been fully established. Model organisms can be used to examine the cellular processes that may be changed by exposure to the E-liquids used for vaping. Mitochondria are essential in eukaryotic cells for production of ATP, protein biogenesis, metabolic pathways, cellular signalling, stress responses and apoptosis. Mitochondrial health can be used as a proxy for many aspects of healthy cellular physiology. Mutations in the PINK1 gene can lead to mitochondria-specific autophagy deficiency. We exposed two strains of *Caenorhabditis elegans*, CB5600 control and CC46 pink1, with 10% concentrations of nine different flavoured E liquids. We measured lifespan, movement, body size, brood size, and we examined their mitochondrial networks to investigate the effect of the E-liquids. We show that the CC46 (pink1) strain is affected by the E-liquids, even the flavours without nicotine, and that they have reduced lifespan, movement ability and mitochondrial organisation. We found that some E-liquids can dramatically shorten lifespan in this strain. Our data emphasise a need to carefully ascertain the potential harm that may be caused by the use of E-liquids.

## Introduction

Electronic cigarettes (EC) and vaping are promoted as a safer alternative to smoking tobacco and have rapidly gained in popularity. This new habit which includes the use of nicotine has seen a rapid increase of users - from ∼7 million in 2011 to 35 million in 2016. According to market research by Euromonitor there are estimates that in adults the numbers of vapers could reach around 55 million by 2021. This rapid increase is also seen in the UK where 6.3% of British adults vaped in 2018, up from 5.5% in the previous year (NHS Digital, July 2019). The most common reason stated for this habit is they are considered “less harmful than regular cigarettes”. Nearly half of regular users said that they vape to reduce smoking tobacco cigarettes. A major concern is that flavoured nicotine vapes are more appealing to children who can be less aware that these are addictive [1]. In 2018, nearly 5% of middle school children and more than 20% of high school children had used vapes in the last month in the USA [2].

When vaping inhaled vapours are generated from an E-liquid containing a mixture of vegetable glycerol (VG), propylene glycol (PG), flavours and nicotine [3]. As they are heated by the electric resistance wire in the EC, VG and PG aerosolise, along with the flavouring and nicotine [4]. This aerosol consists of a suspension mixture of gases, vapours and aqueous particles. The flavourings used in ECs have generally been recognized as safe for use as food additives. However, the consumption of such chemicals has raised concerns due to potential toxicity arising from their inhalation [5]. Although some physiologic effects on *C. elegans* of E-liquids, at a concentration of 0.2%, has been tested [6], there are still very little *in vivo* data available on the safety of different flavourings used for vaping. Mitochondria are vital organelles in eukaryotic cells and their physiological state can be used as a proxy marker for cellular or organism health. These double membraned structures perform important roles, most notably in bioenergetics and metabolic pathways [7],[8], but also in protein biogenesis [9],[10], cell signalling [11], stress response and apoptosis [12]. Mitochondrial dysfunction is associated with ageing and disease [13]. Mutations in the PTEN induced putative kinase 1 (PINK1) gene, which is recruited to damaged mitochondria, can lead to autosomal recessive Parkinson’s disease (PD) [14]. PINK1 and another protein (also associated with familial PD) parkin, act in synergy to reduce a build-up of aged and dysfunctional mitochondria via a process known as mitophagy [15]. Interestingly PINK1 and parkin are also found to be important in handling inflammation [16,17].

The model organism *C. elegans*, as a transparent multicellular eukaryote, has been widely used to study development [18], stress [19], ageing [20] and other biological processes [21]. *C. elegans* with its short lifespan and easy operability is an ideal model organism for the study of environmental toxicology or chronic drug exposure [22]. Initial observations in worms can shed light upon the effects of exposures to basic cellular physiology, leading to more focussed studies in mammalian models and humans [23]. Here we have analysed two strains of *C. elegans*, CB5600 (control) and CC46 (*pink-1* knock out), to investigate the effects of E-liquids on important parameters of physiological health.

## Methods

### General methods and strains

All the E-liquids were re-labelled with numbers, and experiments were conducted blind. E-liquid information is available in Table 1. The concentration of E-liquid: 100ul *E.coli* strain OP50 had 10ul vape liquid added to give a 10:1 ratio. The strains, wild type CB5600 (ccIs4251 I; him-8 (e1489) IV) with GFP fusion proteins localised to nuclei and mitochondria of body wall muscles and CC46 (ccIs4251 I; pink-1(tm1779) II; him-8 (e1489) IV) with GFP fusion proteins localised to nuclei and mitochondria of body wall and muscles were obtained as a generous gift from the laboratory of Prof. Nathaniel Szewczyk. Worm stocks were subsequently maintained at 20°C.

**Table 1.** E-liquids of different flavours and nicotine concentration

| E-liquids | Flavours | Nicotine concentration mg/ml | Brand |
| --- | --- | --- | --- |
| 1 | Vanilla Custard | 3 | Premium Britain E-liquid |
| 2 | Blueberry HVG | 3 | Premium UK E-liquid |
| 3 | Strawberry Milkshake | 3 | Premium UK E-liquid |
| 4 | Blueberry | 3 | Premium UK E-liquid |
| 5 | Very Berry | 0 | Premium UK E-liquid |
| 6 | Heizen* | 3 | Premium UK E-liquid |
| 7 | British Tobacco | 6 | Premium UK E-liquid |
| 8 | Cherry Menthol | 3 | Premium UK E-liquid |
| 9 | Very Berry | 3 | Premium UK E-liquid |

\*Heizen flavour contains mixed fruit, aniseed & menthol (contains anethole & eucalyptol).
All the E-liquids contain vegetable glycerol and propylene glycol.

### Synchronisation of L1 larval animals using gravity flotation

3ml M9 buffer was added to a single plate with high numbers of L1 larvae, and pipetted against the surface of the agar with the plate inclined. The worms in M9 were transferred to a 15 ml tube and washed three times. After leaving to settle for 2.5 min, the top 500 µl of M9 buffer from the tube, containing L1 larvae, was pipetted onto a new NGM plate seeded with 500 µl OP50. The plates were monitored daily and worms transferred on to fresh OP50 NGM plates when the food source had been consumed.

### Life span assay

Upon reaching L4 stage 140 worms of each group were picked out onto nematode growth medium (NGM) agar and fed OP50 with 10% concentration of E-liquid at 20 °C. Worms were transferred daily until the cessation of egg laying to avoid overlapping generations. After the egg laying period finished the worms could be transferred every other day. The worms were scored as dead when they did not respond to touch stimulus.

### Movement assay

The movement assay was conducted every two days. 5 adult worms of each group were picked into 50 µl M9 buffer on a microscope slide. The number of left and rightward body bends in 10 sec were counted, where one leftward and one rightward bend was equal to one stroke. The average from the 5 measurements of the body strokes was taken, then multiplied by six to obtain the movement rate per min.

### Body size measurements of worms

After being exposed to the different E-liquids and OP50 for 4 days at 20°C, 3-5 worms were mounted on a slide (in 20 µl M9 buffer), examined and photographed with a Leica CTR5000. The body lengths were measured from the mouth to the tail tip, while the body widths were measured from side to side at the position of the vulva. These worms were discarded after imaging and not used for further experiments. All length measurements were performed with Image J software [24].

### Imaging of mitochondrial networks in body wall muscle

After being exposed to the different E-liquids and OP50 for 4 days at 20°C, 3-5 worms were picked from the plates into 20 µl M9 on a microscope slide. A cover slip was applied and worms imaged with a Leica CTR5000 under a GFP filter (460∼500 nm, blue light excitation) at 63× and 100× magnification.

### Statistics

All statistical analyses were undertaken using GraphPad Prism (GraphPad Software, Inc., La Jolla, CA, USA).

## Results

### The effects of 9 different E-liquids on *C. elegans* lifespan

We assessed the lifespan of two strains of worms CB5600 (Figure 1a) and CC46 (Figure 1b) by adding 9 different E-liquids made in the UK. The median survival in days exposing different E-liquids to CB5600 were all 5 days. The median survival in days exposing different E-liquids to CC46 varied from 9 days in the control (no E-liquid) to 3 days (*Heizen*). We did not find any significant changes with the E-liquids on strain CB5600. However, all the E-liquids (except for *Very Berry)* shortened the lifespan of CC46 worms compared with the control.

**Figure 1.**
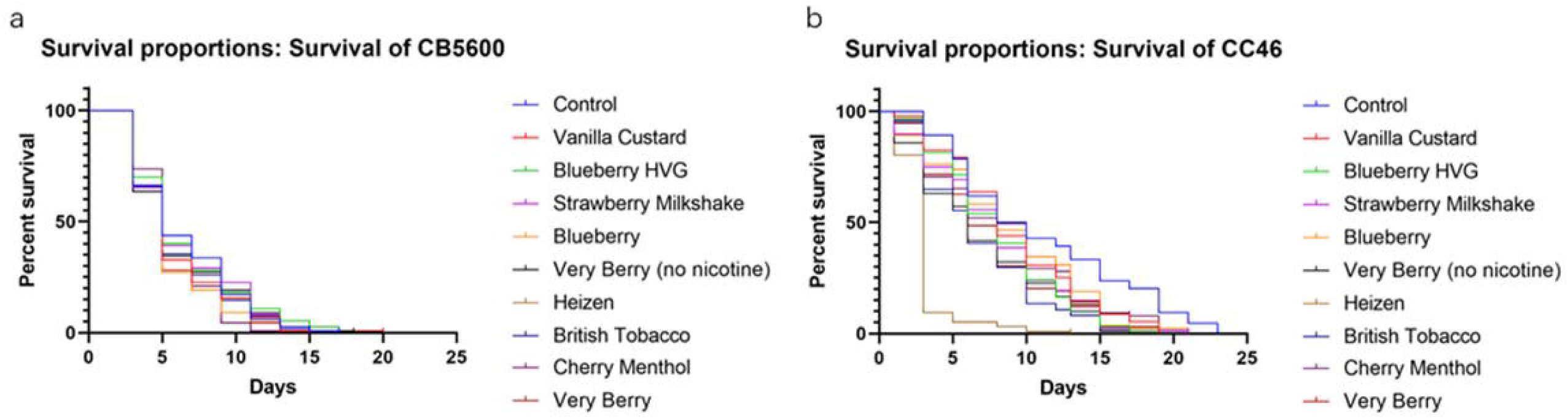
Life-span of *C. elegans* in different E-liquids. Colours represent different E-liquids. Kaplan-Meier graphs show percentage survival. (a) Effect of different E-liquids on lifespan of CB5600. (b) Effect of different E-liquids on lifespan of CC46. The comparison of survival curves was analysed by log-rank (Mantel-Cox) test. We conducted 3 independent tests, ∼ 140 worms in each group were analysed in both CB5600 and CC46.

### Assay to quantify movement defects

Movement assays were carried out every two days after the addition of different E-liquids. Movement declined in all the groups in both CB5600 (Figure 2a) and CC46 (Figure 2b) worms with time. In the older worms there were some significant differences in body movement between the different E-liquids, there were greater differences in the CC46 worms. At 12 days the CB5600 worms exposed to *Cherry Menthol* had a significantly reduced movement rate when compared with the control. CC46 worms exposed to *Blueberry HVG* had a significantly reduced movement rate when compared with controls and the group exposed to the *British Tobacco* E-liquid.

**Figure 2.**
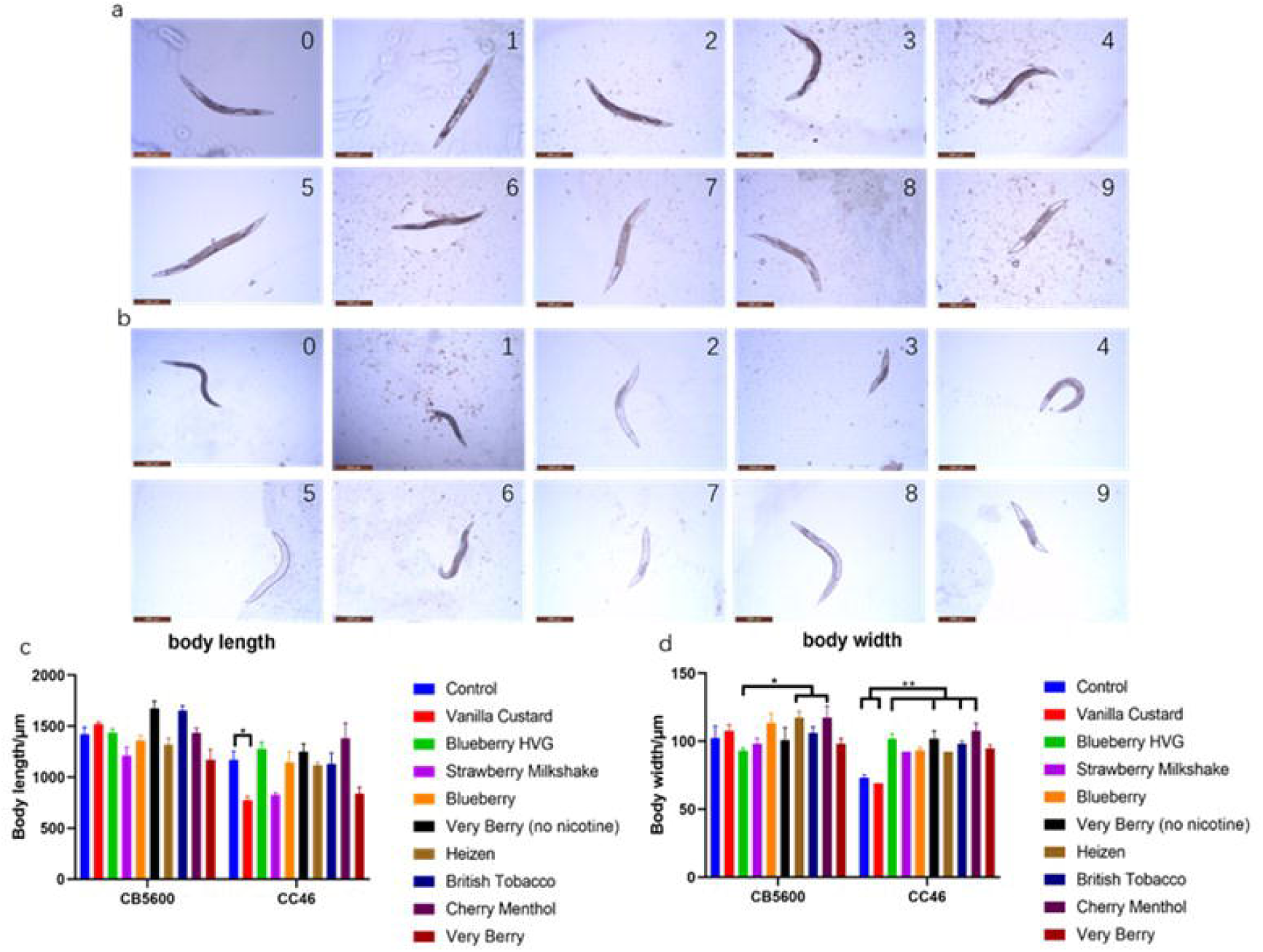
Locomotion of *C. elegans* over time. (a) Movement rate of CB5600 with different E-liquids. (b) Movement rate of CC46 with different E-liquids. Data were analysed using Two-way ANOVA and presented as Mean□±□SEM, multiple comparisons were conducted by Dunnett’s multiple comparisons test. We did not compare E-liquid *Heizen* on the 8-day and 12-day period in CC46, since all the worms were dead at this stage. ^*^ *p*<0.05, ^**^ *p*<0.01.

### Assessment of mitochondrial network structure

To determine the possible underlying mechanisms behind the reduction of movement, we looked at the integrity of the mitochondrial network of both CB5600 and CC46, both strains express GFP in the mitochondria and nuclei of the body wall muscles. Mitochondria provide the bulk of cellular energy, thus the structure of mitochondria were examined for abnormalities. In *C.elegans* CB5600 body wall muscle, there was disruption of the mitochondrial network in all the E-liquid groups (Figure 3a), while in CC46, all the groups also showed fragmentation of the mitochondrial network (Figure 3b).

**Figure 3.**
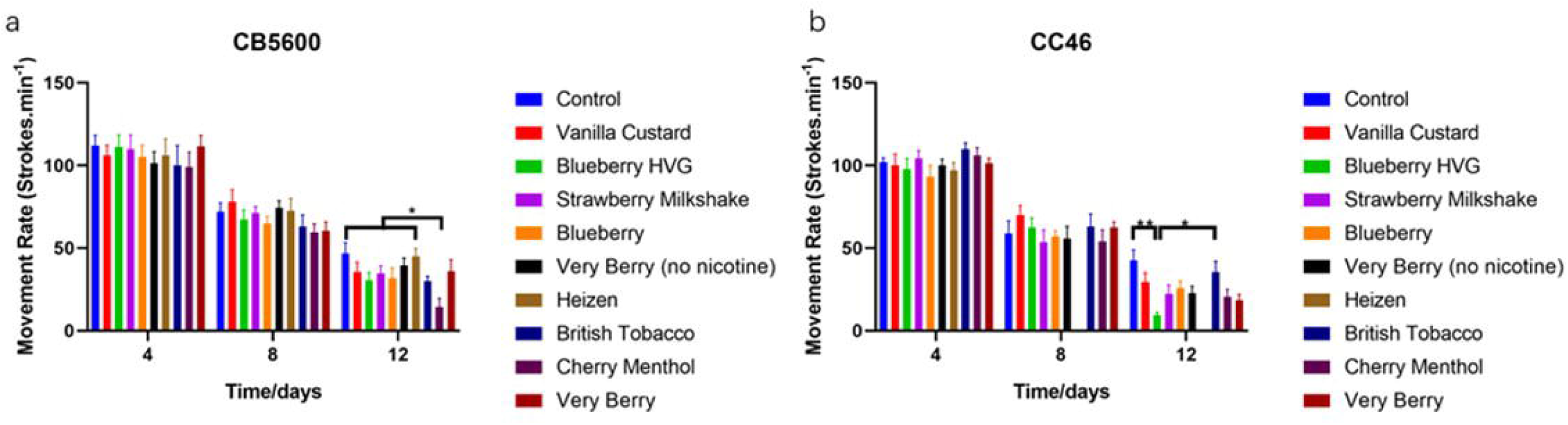
Mitochondrial morphology of *C. elegans*. (a) Mitochondrial morphology of CB5600 after 4 days culture with different E-liquids. (b) The mitochondrial morphology of CC46 after 4 days culture with different E-liquids. Figure numbers represent the different E-liquids: 0, Control; 1, Vanilla Custard; 2, Blueberry HVG; 3, Strawberry Milkshake; 4, Blueberry; 5, Very Berry; 6, Heizen; 7, British Tobacco; 8, Cherry Menthol; 9, Very Berry. The scale bar is 50 µm.

### Body size of worms

Body size was measured as width and length of the worms using imaging software (Figure 4). Significant changes in body length were only found when comparing CC46 controls with *Vanilla Custard* exposed CC46 worms. There appeared to be more of an effect on body width. In CB5600 worms there was a different response (thinner worms) with *Blueberry HVG* when compared with *Heizen* and *Cherry Menthol* (fatter). The CC46 control and *Vanilla Custard* exposed worms were thinner than the others, and significantly more so than CC46 exposed to *Blueberry HVG, Very Berry* (no nicotine), *British Tobacco* and *Cherry Menthol*. The *Cherry Menthol* exposure caused both strains to become significantly fatter.

**Figure 4.**
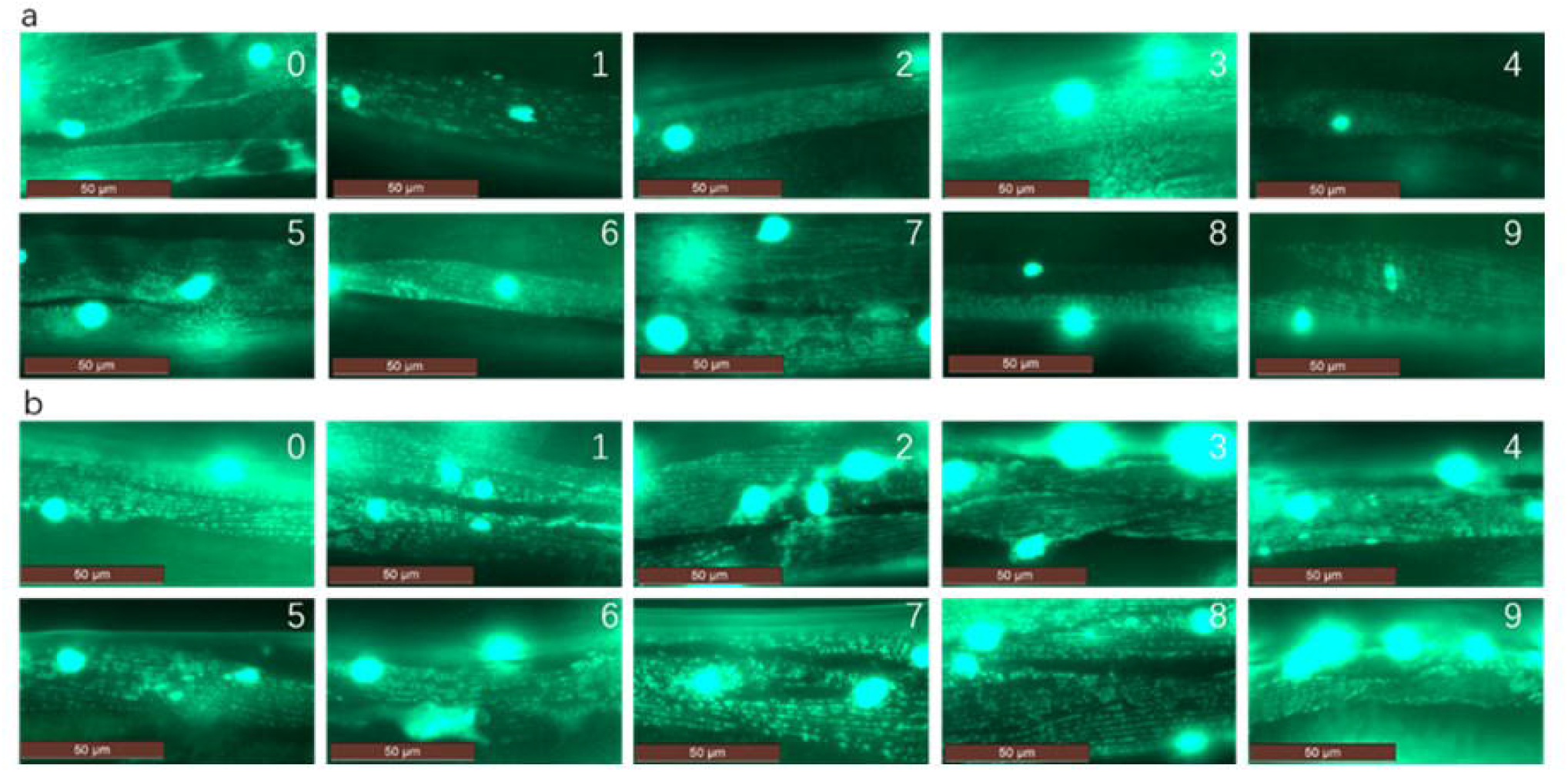
The body size of *C. elegans*. (a) The morphology of CB5600 after 4 days culture with different E-liquids. (b) The morphology of CC46 after 4 days culture with different E-liquids. The scale bar is 500µm. (c) Body length of CB5600 and CC46 after 4 days culture with different E-liquids. (d) Body width of CB5600 and CC46 after 4 days culture with different E-liquids. Data were analysed using One-way ANOVA and presented as Mean□±□SEM, multiple comparisons were conducted by Tukey’s multiple comparisons test. ^*^ *p*<0.05, ^**^ *p*<0.01.

## Discussion

The current view that the use of E-cigarettes and ‘vaping’ is relatively safe is based on little evidence. Initial studies were largely epidemiological to examine the take-up of this habit and dependency in vulnerable and young people [25]. Now there is a growing body of evidence outlining potential harmful effects on inhaling E-liquid vapours [26].

We set out to examine the effects of different flavour vapes on the lifespan, movement, body size, brood size, and mitochondrial networks in two strains of *C. elegans*. The physiological consequences of exposing *C. elegans* to different vapes can give us important information regarding the potential harm that may be caused by the use of E-liquids.

We chose strains that might shed light on the effect of vapes in organisms with metabolic changes affecting mitochondrial physiology. The CB5600 were the control strain against which we measured CC46, a strain with deficiency in PINK1, a protein involved in mitophagy and recently implicated in inflammation [16]. Lifespan in the CC46 worms was longer than the CB5600 overall. However, the effect of E-liquid exposure on the PINK1 deficient strain resulted in a greater reduction of their expected lifespan. We included one E-liquid that didn’t contain nicotine. Nicotine has been tested in worms and demonstrated to cause muscle paralysis, affect egg laying and also spicule ejection [27]. Higher nicotine concentrations (20 to 30 mM in liquid culture), cause rapid (within 10-15 minutes) paralysis of body wall muscle in wild-type young adults. If allowed a recovery period of 45-60 min after exposure the worms can acquire tolerance [27]. Lower nicotine concentrations (6.17–194.5 μM) may also affect stimulus-response, reproductive function and gene expression of *C. elegans* [28]. In our study, all the E-liquids except for one contained nicotine, but the nicotine final concentration was very low (0.1μM). The *Very Berry* flavour without nicotine didn’t exhibit any survival advantage over the worms treated with nicotine containing liquids. When control CB5600 worms were exposed to E-liquid we did not see substantial deaths in a short time with the products, thus we can conclude that in this study nicotine is not a strong factor influencing worm death. The largest reduction in lifespan was with the E-liquid *Heizen*, which was different to the other vapes in containing aniseed and menthol (anethole and eucalyptol), these ingredients may have induced the acute reaction in CC46 worms. Previous research has showed that eucalyptol has toxic effects on MRC-5, HT-29 and HCT 116 cells and also may result in maternal and fetal toxicity in the rat [29],[30]. We suggest that the eucalyptol in this E-liquid may have highly toxic effects in cells and tissues where mitochondrial physiology is already compromised.

Movement assays are a good measure of functional physiology in the nematode worm. As might be anticipated the frequency of movement declined with age. *Heizen* exposed CC46 worms did not survive through to the second and third time points but performed as well as the others and controls in the earliest timepoint. Again, *Cherry Menthol* appears to affect the CB5600 worms more than any of the other flavours. Menthol has been investigated for various properties such as muscle or airway relaxation and therefore it may be these that are reducing the movement of the worms [31]. In the context of E-liquids menthol has been found to increase neonatal pulmonary artery smooth muscle cell death, suggesting that there may be good reason to restrict menthol use in order to protect lungs of young vapers and also guard against pre-natal exposures [31].

Muscle mitochondria are distributed in evenly dispersed networks which can be readily visualised with GFP in the CB5600 and CC46 strains we used. Representative images show that many of the E-liquids are associated with disruption of the networks. All the E-liquids disrupted the mitochondrial networks and the extent of the disruption is greater in the CC46 strain which already has a mitochondrial deficiency. Importantly E-liquid *Very Cherry* without nicotine (Figure 3b-5) shows large disruptions, with clumps and gaps of GFP fluorescence, and *British Tobacco* (Figure 3b-7) which has twice the quantity of nicotine as the rest is also strongly affected. This again demonstrates that the effects of nicotine may not be the most important. Mitochondrial networks were imaged when the worms were younger to control for the divergent lifespans in the two strains. Mitochondrial function is key to survival for obligatory aerobic eukaryotes, this is confirmed by the toxic effect of molecules that interfere with their integrity [32]. The nicotine concentrations in our study were very low and might be most comparable to the levels found in the environment when the cigarette has stopped burning, called third-hand cigarette smoke (THS). It has been shown that THS from just a few cigarettes can effect stress responses in mitochondria and change the transcriptional profile of genes associated with mitochondrial function, this must affect the mitochondria and implicitly also affect cellular health [33]. If exposure to THS is prolonged then mitochondrial membrane potential (MMP) and cell proliferation decrease [34]. A low dose of THS may not be directly responsible for cell death but the stressed state it induces compromises cellular function, survival and proliferation, potentially leaving individuals more vulnerable to ill-health [35].

Body length and width were measured after four days of exposure to the E-liquids. Body length did not significantly change except with *Vanilla Custard* in CC46. There do seem to be other flavours that probably affect the CC46 body length though not sufficiently to be recorded as significant, the changes mostly cause the worms to be shorter. Body width appears to be more generally affected in the CC46 strain with many of the groups getting fatter than the controls. There are investigations to probe the relationship of weight and vaping habits as a corollary to knowledge that smoking tobacco can reduce or help maintain body weight [36]. The studies to establish any causative relationship between weight and E-cigarette have not been done but preliminary work identifies a link between obesity and higher likelihood of E-cigarette use, suggesting that there may be some effect [37].

The CC46 strain overall showed greater sensitivity to the vapes with the tests we used. Mutations in mitochondrial DNA and in PINK1appear to be more frequent in mice exposed to nicotine containing e-cigarettes [38]. There is recent work connecting PINK1 and mitochondrial stress with the disruption to innate immunity and the STING pathway and this if conserved in *C.elegans*, may provide some explanation for the effects we see [16][39]. Indeed, there is already evidence connecting disruption of immunity mechanisms with the use of E-liquids where the effect of nicotine is laso suggested to not be the most important [40][41]. The CC46 model could be useful for further studies into conserved pathways that are affected by using E-liquids.

Both *Cherry menthol* and *Heizen* flavours showed consistent detrimental effects on the strains throughout this study. Eucalyptol is the ingredient that is specific to the *Heizen* flavour and has recently been demonstrated to be useful as a therapy for lung damage after cigarette smoke exposure [42–44]. Eucalyptol, also known as 1,8-cineole has been used to flavour food and therapeutically in nasal-sprays, muscle creams and has been associated with anti-inflammatory properties. Astonishingly, information about its action at the cellular or sub-cellular level has not been clearly defined, sub-acute levels have been shown to affect liver and kidney tissues causing haemorrhagia, swelling and granular degeneration [45]. Ultrastructural and respirometry changes were noted when evaluating the mitochondrial component of these tissues. Menthol and eucalyptol have been investigated in human endothelial cells the context of cardiovascular health and found to potentially disrupt nitric oxide production pathways [46]. Mentholated cigarettes are banned in Europe and a ban is also being explored by the FDA. The state of Massachusetts temporarily restricted the sale of all menthol and flavoured cigarette and vape products following some acute reactions to these [47]. The ban has now been lifted.

There is concern for various reasons about the use and unrestricted sale of E-cigarettes, particularly those containing menthol. We find that *C.elegans* physiology is clearly affected by some E-liquids we tested. The effect upon worms with a known genetic deficiency affecting mitochondrial pathways is amplified, perhaps indicating that the baseline metabolic status of the individual who is using these substances could be very relevant for E-liquid safety profile. It is important now to delineate the precise effects of E-liquids and their components on cellular physiology in order to inform policymakers of their potential to cause harm.

## Author contributions

YW performed the laboratory works of lifespan, movement, body size, brood size, mitochondrial imaging assays, data analysis and manuscript preparation; TLI performed the laboratory works of mitochondrial imaging, data analysis and manuscript preparation; SM performed preliminary laboratory work to determine the vape concentration; FS performed some preparation of worm culture laboratory work; LC conceived the study, provided reagents and prepared the manuscript.

## Supplementary data

**Table 1.** The significance of body length comparisons between all the groups in CB5600

| Groups | *p value* | Summary |
| --- | --- | --- |
| Vanilla Custard vs. Strawberry Milkshake | 0.0227 | * |
| Vanilla Custard vs. Very Berry | 0.0117 | * |
| Strawberry Milkshake vs. Very Berry (no nicotine) | <0.0001 | **** |
| Strawberry Milkshake vs. British Tobacco | 0.0001 | *** |
| Blueberry vs. Very Berry (no nicotine) | 0.0185 | * |
| Blueberry vs. British Tobacco | 0.0372 | * |
| Very Berry (no nicotine) vs. Heizen | 0.0092 | ** |
| Very Berry (no nicotine) vs. Very Berry | <0.0001 | **** |

**Table 2.** The significance of body length comparisons between all the groups in CC46

| Groups | *p value* | Summary |
| --- | --- | --- |
| Vanilla Custard vs. Blueberry HVG | 0.0010 | *** |
| Vanilla Custard vs. Blueberry | 0.0426 | * |
| Vanilla Custard vs. Very Berry (no nicotine) | 0.0023 | ** |
| Vanilla Custard vs. Cherry Menthol | <0.0001 | **** |
| Blueberry HVG vs. Strawberry Milkshake | 0.0045 | ** |
| Blueberry HVG vs. Very Berry | 0.0070 | ** |
| Strawberry Milkshake vs. Very Berry (no nicotine) | 0.0100 | ** |
| Strawberry Milkshake vs. Cherry Menthol | 0.0002 | *** |
| Very Berry (no nicotine) vs. Very Berry | 0.0151 | * |
| Cherry Menthol vs. Very Berry | 0.0003 | *** |

